# 3D Echo Planar Time-resolved Imaging (3D-EPTI) for ultrafast multi-parametric quantitative MRI

**DOI:** 10.1101/2021.05.06.443040

**Authors:** Fuyixue Wang, Zijing Dong, Timothy G. Reese, Bruce Rosen, Lawrence L. Wald, Kawin Setsompop

## Abstract

Multi-parametric quantitative MRI has shown great potential to improve the sensitivity and specificity of clinical diagnosis and to enhance our understanding of complex brain processes, but suffers from long scan time especially at high spatial resolution. To address this long-standing challenge, we introduce a novel approach, termed 3D Echo Planar Time-resolved Imaging (3D-EPTI), which significantly increases the acceleration capacity of MRI sampling, and provides high acquisition efficiency for multi-parametric MRI. This is achieved by exploiting the spatiotemporal correlation of MRI data at multiple timescales through new encoding strategies within and between efficient continuous readouts. Specifically, an optimized spatiotemporal CAIPI encoding within the readouts combined with a radial-block sampling strategy across the readouts enables an acceleration rate of 800 folds in the *k-t* space. A subspace reconstruction was employed to resolve thousands of artifact-free high-quality multi-contrast images spaced at a time interval of ~1 ms. We have demonstrated the ability of 3D-EPTI to provide robust and repeatable whole-brain simultaneous T_1_, T_2_, T_2_*, PD and B_1_^+^ mapping at high isotropic resolution within minutes (e.g., 1-mm isotropic resolution in 3 minutes), and to enable submillimeter multi-parametric imaging to study detailed brain structures.

**Highlights:**

- Ultra-fast acquisition for 3D multi-parametric quantitative MRI.
- Simultaneous T_1_ T_2_ T_2_* PD and B_1_^+^ mapping.
- 3-minute scan at 1-mm isotropic resolution with whole-brain coverage.
- Multi-parametric mapping at 700-μm isotropic resolution in 10 minutes.
- Repeatable quantification and cortical-depth analysis.

## 1. Introduction

Multiparametric MRI provides quantitative measurements that are sensitive to a variety of tissue properties of the human brain. Its quantitative nature leads to less reliance on system conditions and human interpretation compared to standard qualitative images, and makes it possible to measure tissue properties among populations or along time, therefore providing great potentials to improve clinical diagnosis (Bernasconi et al., 1999; Falangola et al., 2007; Lescher et al., 2015; Ma et al., 2018; Müller et al., 2017; Ramani et al., 2006; Reitz et al., 2017; Tardif et al., 2011; Tofts, 2005; West et al., 2014) or to enhance our understanding of complex brain processes such as brain development or aging (Bozzali et al., 2016; Filo et al., 2019; Sled and Nossin-Manor, 2013).

A major limitation of quantitative multiparametric MRI is its long acquisition time. This is due to the need to acquire a series of multi-contrast images in order to perform model fittings for quantitative parameter estimation, as well as the need to repeat such process for each parameter in multiparametric imaging. Acceleration of the acquisition has been made possible by using parallel imaging (Griswold et al., 2002; Pruessmann et al., 1999; Sodickson and Manning, 1997) or by enforcing prior knowledge of image properties through compressed-sensing theory (Lustig et al., 2007), both of which can help recover undersampled *k*-space. Together with new pulse sequence design, many fast quantification methods have been developed to obtain multiple parameters simultaneously (e.g., T_1_, T_2_, T_2_*) (Caan et al., 2019; Deoni et al., 2003; Fujita et al., 2019; Krauss et al., 2015; Metere et al., 2017; Warntjes et al., 2007; Warntjes et al., 2008). However, the ability to accelerate the acquisition is still limited and obtaining high-resolution multi-parametric MRI in clinical acceptable time remains a challenge.

Recently, an emerging area of research has been in the use of spatiotemporal correlation to achieve high acceleration for quantitative MRI. For example, MR fingerprinting (Boyacioglu et al., 2021; Cloos et al., 2016; Hong et al., 2019; Jiang et al., 2015; Ma et al., 2013; Ma et al., 2018; Wyatt et al., 2018) and MR multitasking (Christodoulou et al., 2018; Ma et al., 2020; Ma et al., 2021), have utilized spatiotemporal correlation between readouts after different RF excitations to accelerate multi-parametric imaging, which have shown promising results especially when used in conjunction with the low-rank subspace model (Christodoulou et al., 2018; Liang, 2007; Zhao et al., 2018). Another approach, Echo Planar Time-resolved Imaging (EPTI) (Dong et al., 2021a; Dong et al., 2020; Wang et al., 2019a; Wang et al., 2019b; Wang et al., 2021), has been developed recently to exploit a stronger spatiotemporal correlation within a continuous EPI-like readout. Within the readout, data are sampled at a much shorter timescale (submillisecond), so only minimal phase accumulation and signal decay will occur, resulting in a stronger temporal correlation that can increase the ability to recover highly undersampled data. In addition, it better takes advantage of the spatial information from multi-channel coils by employing a novel spatiotemporal encoding strategy within the readout, therefore allowing high acceleration. The continuous EPTI readout enables high sampling efficiency with minimal dead time, while the time-resolving approach across the readout eliminates the undesirable image distortion and blurring common in the conventional EPI (Mansfield, 1977), providing a series of high quality multi-contrast images sampled at small time interval (~1 ms) that can continuously track the signal evolution to fit quantitative parameters. We have demonstrated efficient whole brain T_2_ and T_2_* mapping using a GE-SE EPTI at 1.1-mm inplane resolution within 28s in our previous work (Wang et al., 2019a).

Here, in pursuit of a significant further increase in acceleration capability, 3D-EPTI has been developed. 3D-EPTI extends the spatiotemporal EPTI encoding from 2D (*k_y_-t*) to 3D (*k_y_-k_z_-t*), and develops a new data sampling strategy with a combined controlled- and incoherent-aliasing scheme. Data correlations at multiple timescales are exploited, both within and between the continuous readouts. Within readouts, a spatiotemporal encoding is designed with complementary sampling in a controlled-aliasing (Breuer et al., 2005; Breuer et al., 2006; Dong et al., 2019) (CAIPI) pattern in the *k_y_-k_z_-t* domain, which uses coil sensitivity information along both partition and phase encoding directions along with the temporal correction across echoes, and offers a remarkably higher acceleration capacity (e.g., 80×). Between readouts, a novel radial-block encoding is developed to exploit data correlation in a longer timescale based on the compressed sensing theory (Lustig et al., 2007). The radial-block sampling creates incoherent aliasing along time that can be well excluded from the coherent signal evolutions, therefore provides another 10× acceleration. The combination of spatiotemporal CAIPI and radial-block undersampling offers a remarkable ~800× acceleration in the spatiotemporal domain.

The harmonious integration of the continuous readout and the novel intra- and inter-readout spatiotemporal encoding in 3D-EPTI provides unique datasets that allow us to time-resolve thousands of multi-contrast 3D images by enforcing the spatiotemporal correlation in the reconstruction process. While this acquisition scheme can be applied or adapted to any sequence or other types of readouts to accurately track the signal evolution and to measure a variety of quantitative parameters, this work employs specific sequence to simultaneously obtain MR relaxation time constants T_1_, T_2_, T_2_* as well as RF field inhomogeneity (B_1_^+^) and proton density (PD). We demonstrated the ability of 3D-EPTI to acquire high-quality whole-brain multi-parametric maps at isotropic 1.5-mm in 1 minute or at isotropic 1-mm in 3 minutes with high repeatability and reliability. An isotropic 0.7-mm 3D-EPTI protocol was also developed to allow, for the first time, the examination of simultaneously acquired T_1_, T_2_, and T_2_* in less than 10 minutes to help better investigate the intra-cortical architecture.

## 2. Material and Methods

### 2.1. 3D-EPTI acquisition

Figure 1 illustrates the 3D-EPTI acquisition. An inversion-recovery gradient echo (IR-GE) and a variable-flip-angle gradient-and-spin-echo (Feinberg and Oshio, 1991; Oshio and Feinberg, 1991) (VFA-GRASE) sequence (Fig. 1a) were chosen and carefully optimized to provide signal evolutions that are sensitive to T_1_, T_2_, and T_2_* relaxations (Fig. 1b). After each RF excitation, a 3D-EPTI readout is acquired, which continuously captures the temporal signal evolution with efficient bipolar gradient. To resolve images within the readout with high acceleration, a spatiotemporal CAIPI encoding is employed in a 4D spatiotemporal (*k_x_-k_y_-k_z_-t*) domain (Fig. 1d, the readout dimension *k_x_* is fully-sampled and therefore omitted in the illustration). At each time point within the readout, a particular phase and partition position (*k_y_-k_z_*) is acquired that is interleaved to its neighboring time points in a ‘controlled-aliasing’ pattern. Across a slightly longer timescale, two complementary CAIPI patterns are interleaved across echoes (orange and green points in Fig. 1d, different ‘echo sections’) to provide more independent *k*-space sampling locations, which has been shown to further improve the reconstruction performance at high acceleration rates (Dong et al., 2020). Each 3D-EPTI readout covers a relatively small block in *k_y_-k_z_-t* space to ensure that the neighboring *k_y_-k_z_* samplings are close in time. The CAIPI pattern, the complementary sampling across echoes, and the proximity in time together result in high spatiotemporal correlation and allow effective use of the available coil sensitivity information. Therefore, the highly-undersampled data (e.g., undersampling rate = 80×) at each time point can be well reconstructed, resolving a series images across the readout at a submillisecond time interval (Fig. 1d, left).

**Figure 1.**
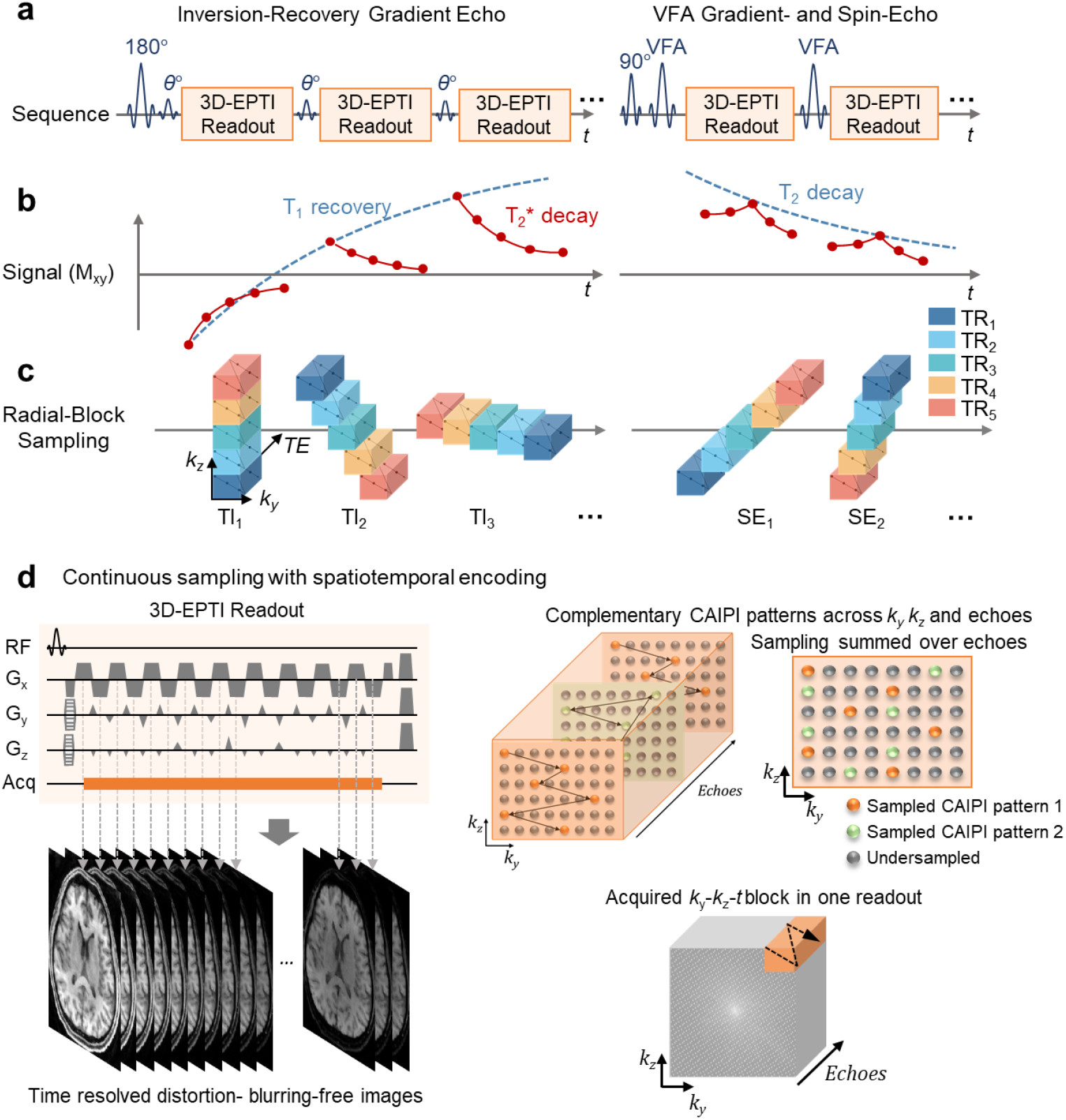
Illustration of the 3D-EPTI acquisition. **a**, The sequence diagrams of the inversion-recovery gradient-echo (IR-GE) and the variable-flip-angle gradient and spin-echo (VFA-GRASE) sequences with 3D-EPTI readouts at different inversion times (TI) and spin-echoes (SE). **b**, The designed sequence provides signal evolutions with high sensitivity to T_1_, T_2_ and T_2_* relaxation time constants, which can be continuously tracked by the 3D-EPTI readouts. **c**, Instead of acquiring the full *k_y_-k_z_* space at every TI or SE, a radial-block Cartesian sampling pattern is utilized to quickly sample the *k-t* space in a small number of TRs. At each TI or SE, the *k_y_-k_z_-t* blocks acquired in different TRs (color-coded) form a radial blade with different angulations to create spatiotemporal incoherent aliasing for constrained reconstruction and permit ~10× acceleration. **d**, Details of the continuous bipolar readout with an optimized spatiotemporal CAIPI encoding used to efficiently cover a *k_y_-k_z_-t* block per 3D-EPTI readout. The neighboring data points are acquired close in time to create strong temporal correlation. Two CAIPI patterns (orange and green points) are utilized in a complementary fashion at a longer timescale in different echo sections. The combination of spatiotemporal CAIPI and radial-block undersampling offers a remarkable ~800× acceleration in the spatiotemporal domain. Note that the readout (*k*_x_) dimension is fully-sampled and therefore omitted in the illustration.

In each repetition time (TR), multiple *k-t* blocks can be acquired across multiple readouts after different excitations (Fig. 1c, blocks in the same color are acquired in the same TR). To quickly encode the 4D *k-t* space using a small number of TRs, a golden-angle radial-block Cartesian sampling is employed across the readouts. Specifically, the blocks acquired after the same excitation in different TRs form a diagonal radial blade in the *k_y_-k_z_* space, with different blade angulations across different readouts. This was developed to create a favorable spatiotemporal incoherent aliasing across the readouts that is well suited for constrained reconstruction, which permits a further ~10× acceleration through acquiring only a few blades for each readout instead of the full *k_y_-k_z_* sampling.

### 2.2. Image reconstruction

The acquired highly undersampled data, with carefully designed spatiotemporal encoding patterns, will then be reconstructed by a low-rank subspace reconstruction (Dong et al., 2020; Guo et al., 2021; He et al., 2016; Lam and Liang, 2014; Liang, 2007; Meng et al., 2021; Tamir et al., 2017; Zhao et al., 2015), tailored specifically to 3D-EPTI to time-resolve thousands of multi-contrast images (Fig. 2). The low-rank subspace method was chosen for use based on its superior ability to improve the conditioning in the reconstruction by utilizing the low-rank prior information of the signal evolution, therefore achieving high image SNR.

**Figure 2.**
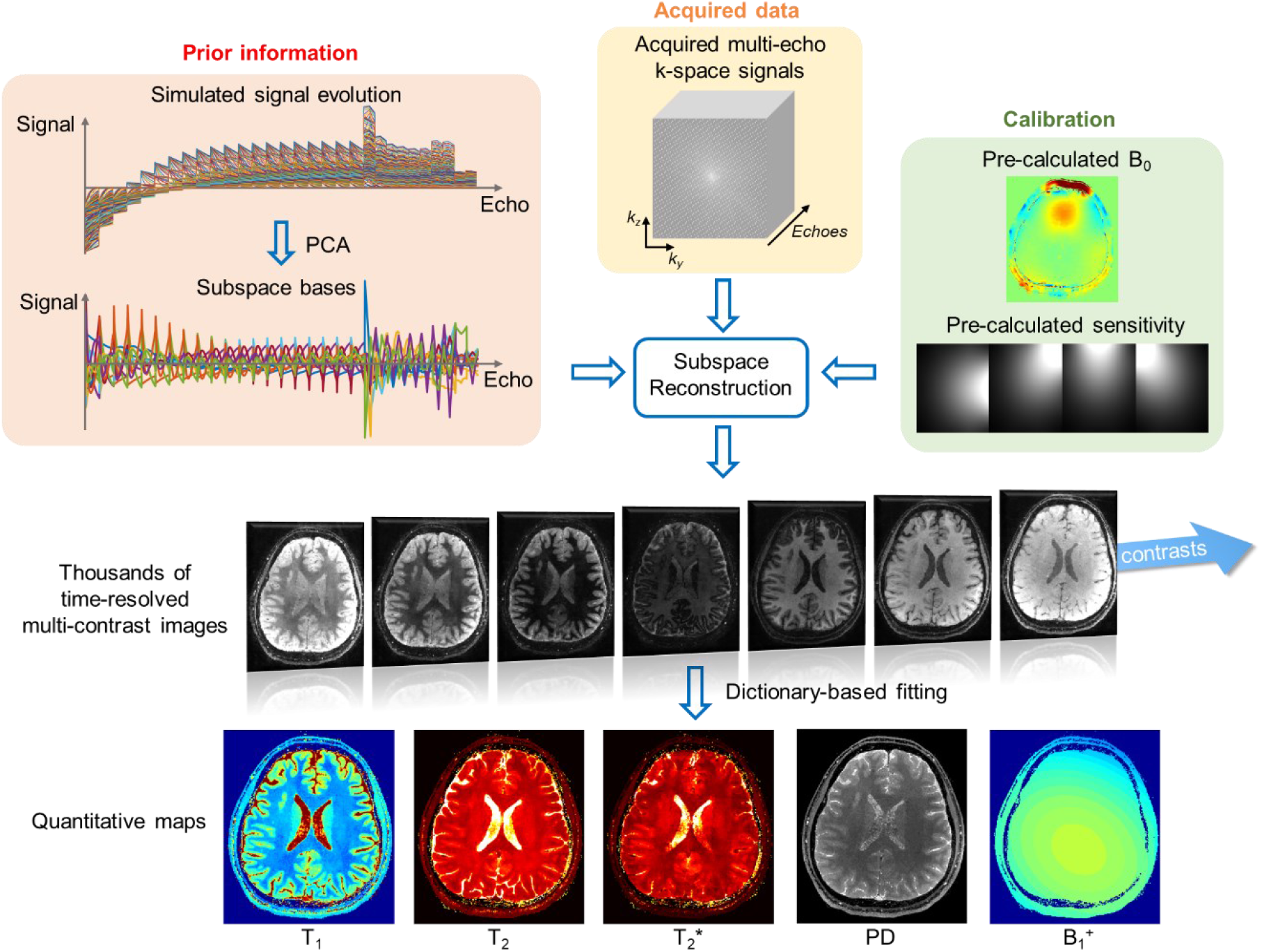
Illustration of the reconstruction framework of 3D-EPTI using the low-rank subspace method. The signal evolutions can be represented by a linear combination of several subspace temporal bases, therefore reducing the number of unknowns from thousands of images to a few coefficients of the subspace bases. The bases are extracted from the simulated signal space using principal component analysis (PCA). The subspace reconstruction is performed by integrating the information from a highly-undersampled spatiotemporal dataset, *B*_0_ phase evolution and coil sensitivity maps obtained via a calibration dataset, and the subspace bases. After the reconstruction, thousands of multi-contrast images can be obtained without distortion and blurring, from which multiple quantitative parameters including T_1_, T_2_, T_2_*, PD, and B_1_^+^ can be estimated through dictionary matching.

At first, a large number of temporal signal evolutions are simulated using the Extended Phase Graphs (EPG) approach (Weigel, 2015), each contains *N_t_* time points. A wide range of quantitative parameters were used to exhaust all possibilities of interest (e.g., T_1_: 400 ms to 5000 ms, T_2_: 10 ms to 500 ms, T_2_*: 10 ms to 500 ms, B_1_^+^ factor: 0.75 to 1.25). Second, *N_b_* subspace basis vectors 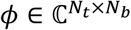 are extracted from these simulated signals by using principal component analysis (PCA). In this study, 12 bases were selected that can approximate the simulated signals with an error smaller than 0.2%. Then, the full time series of *N_ν_* spatial voxels can be represented by *φc*, where 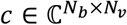 represents the coefficient maps of the subspace bases that can be estimated by solving:

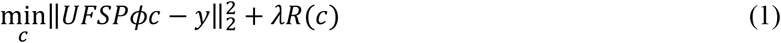

where 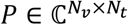 contains the phase evolutions across the time-series images including the background and *B*_0_ inhomogeneity-induced phases, 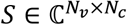 denotes the coil sensitivity of a *N_c_*-channel receiver coil, *F* is the Fourier transform operator, *U* is the undersampling mask, and *y* represents the acquired *k*-space data. The phase map *P* and sensitivity map S’ can be obtained from a short low-resolution calibration pre-scan. A locally low-rank (LLR) regularization *R*(*c*) is applied on coefficient maps with a control parameter *λ* to further improve the conditioning. Since the number of unknowns in the optimization problem is significantly reduced from thousands of images to a few coefficient maps, the subspace method can achieve accurate image reconstruction for 3D-EPTI at high accelerations.

After reconstruction, the quantitative values can be obtained by matching the signal evolution with a pre-calculated dictionary. The dictionary is generated using the EPG approach with the same parameter range used in the basis generation. After dictionary matching, the estimated B_1_^+^ maps are fitted by a 2^nd^-order polynomial function in the 3D spatial domain, with an assumption that B_1_^+^ fields vary smoothly in the spatial domain, which are then fed back into the dictionary matching to obtain more accurate quantitative parameters.

### 2.3. In-vivo experiments and acquisition parameters

All data were acquired on a Siemens Prisma 3T scanner with a 32-channel head coil (Siemens Healthineers, Erlangen, Germany). Informed consent was collected from all healthy volunteers before scanning, with an institutionally approved protocol.

The details of the pulse sequence used in our experiments are described here. In the IR-GE sequence, an adiabatic inversion pulse was applied, followed by 20 excitation pulses with a small flip angle (FA) of 30°. In the VFA-GRASE sequence, 10 variable-flip-angle refocusing pulses were applied after the 90° pulse with FAs of 122°, 58°, 44°, 41°, 41°, 46°, 158°, 189°, 43°, 30°. The FAs and the timing of the pulse sequence were chosen based on the results of an optimization considering both the signal amplitude and the differentiability between tissues. All the excitation and refocusing RF pulses in the IR-GE and GRASE sequences were non-selective with short pulse durations (0.5 ms for excitation and 1 ms for refocusing), resulting in shorter achievable starting TE and sampling interval. Readout gradient was applied along the Head-Foot (HF) direction to avoid signal wrap from the non-selective excitation. Spectrally-selective fat saturation was applied before excitation.

To evaluate the repeatability of 3D-EPTI, a scan-rescan assessment was performed on 5 healthy volunteers using a 3-minute 1-mm 3D-EPTI protocol, where the subjects were taken out of the scanner and repositioned between the two scans. The data were acquired with the following parameters: FOV = 220 × 176 × 210 mm^3^ (AP-LR-HF), matrix size = 230 × 184 × 210, spatial resolution = 0.96 × 0.96 × 1 mm^3^, echo spacing = 0.93 ms, TR of IR-GE = 2600 ms, TR of GRASE = 800 ms, block size (*k_y_* × *k_z_*) = 8 × 10 (80× acceleration, see Supplementary Fig. 1a for encoding pattern). There were 20 readouts in IR-GE, each containing 48 echoes, and 10 readouts in GRASE, each containing 39 echoes. 2 radial lines were acquired for the IR-GE sequence in 45 TRs (instead of 23×23=529 TRs without radial-block acceleration), and 3 radial lines in 65 TRs were acquired for the GRASE sequence (Supplementary Fig. 2 top row). More lines were acquired in GRASE to compensate for the fewer number of readouts to encode the *k-t* space. The total acquisition time was ~3 minutes, including 117 seconds for IR-GE, 54 seconds for GRASE, and a 12-second calibration scan. The *k-t* calibration scan was acquired to estimate the *B*_0_ and coil sensitivity using a GE sequence with bipolar readouts with the following parameters: matrix size = 42 × 32 × 210, number of echoes = 9, TR = 24 ms. The *k*-space center (8 × 8) was fully-sampled and the rest of *k*-space was undersampled along *k_y_* and *k_z_* by a factor of 2 × 2. GRAPPA (Griswold et al., 2002) was used to reconstruct the missing data points in the calibration data. The calibration scan for all of the following acquisitions used the same matrix size along *k_y_* and *k_z_* with the same acceleration factor.

Two additional whole-brain protocols at different resolutions were also acquired. i) A fast 1-min 1.5-mm protocol with parameters: FOV = 218 × 173 × 230 mm^3^ (AP-LR-HF), matrix size = 150 × 120 × 154, spatial resolution = 1.45 × 1.44 × 1.49 mm^3^, echo spacing = 0.72 ms, TR of IR-GE = 1900 ms, TR of GRASE = 600 ms, block size (*k_y_* × *k_z_*) = 8 × 10 (80× acceleration, see Supplementary Fig. 1a for encoding pattern), each readout contained 53 echoes in IR-GE, and 39 echoes in GRASE. 3 radial lines with a reduction factor of 2 along the radial direction, equivalent to 1.5 lines, were acquired in 23 TRs (instead of 15×15=225 TRs), and 2 radial lines were acquired for GRASE in 29 TRs (Supplementary Fig. 2 middle row). The total acquisition time was ~ 1 minute, including 44 s for IR-GE and 17 s for GRASE. The 1.5-mm calibration scan took 10 seconds with a TR = 20 ms. ii) A high resolution 9-min 0.7-mm protocol with acquisition parameters: FOV = 224 × 176.4 × 224 mm^3^ (AP-LR-HF), matrix size = 328 × 246 × 322, spatial resolution = 0.68 × 0.72 × 0.70 mm^3^, echo spacing = 1.2 ms, TR of IR-GE = 2600 ms, TR of GRASE = 800 ms, block size (*k_y_* × *k_z_*) = 8 × 6 (48× acceleration, see Supplementary Fig. 1b for encoding pattern), each readout contained 42 echoes in IR-GE, and 33 echoes in GRASE. 4 radial lines were acquired for both IR-GE and GRASE in a total of 161 TRs (instead of 41×41=1681 TRs) to provide sufficient sampling for higher spatial resolution (Supplementary Fig. 2 bottom row). The total acquisition time was ~ 9 minutes, including 7 min for IR-GE and 2 min for GRASE. The calibration scan took 12 seconds with 7 echoes and a TR of 24 ms.

To further validate 3D-EPTI’s reliability, a comparison study between quantitative parameters provided by 3D-EPTI and those from lengthy standard acquisition methods was conducted both in phantom and *in vivo.* The standard acquisition includes a 2D IR-SE sequence for T_1_ mapping, a 2D single-echo SE sequence for T_2_ mapping, and a 3D multi-echo GRE sequence for T_2_* mapping. In the phantom experiment, the acquisition parameters of the 2D IR-SE sequence were: FOV = 256 × 256 mm^2^, in-plane resolution = 1 × 1 mm^2^, slice thickness = 3 mm, number of slices = 9, acceleration factor along *k_y_* = 2, TR = 8000 ms, TIs = 100, 200, 400, 800, 1600, 3200 ms. The 2D single-echo SE sequence acquired 6 echo times (25, 50, 75, 100, 150, 200 ms) with a TR of 3000 ms, and used the same FOV, resolution, and acceleration as the IR-SE. The 3D multi-echo GRE sequence used for the T_2_* mapping was acquired with a FOV of 186 × 176 × 224 mm^3^ at 1-mm isotropic resolution. Seven echo times (5, 10, 15, 25, 35, 45, 60 ms) were acquired with an acceleration factor of 2 × 2 (*k_y_* × *k_z_*). The total acquisition time for the phantom scan was about 1.5 hours, including 30 minutes for IR-GE, 50 minutes for single-echo SE, and 12 minutes for 3D GRE. The same sequences were used for the in-vivo test, but at a lower resolution of 2 × 2 mm^2^ in the IR-SE and single-echo SE sequences to reduce the scan time and mitigate potential motion-induced artifacts. Most of the other parameters were kept the same as the phantom scan, except for: the number of slices = 19 for IR-SE and single-echo SE; TEs of single-echo SE = 25, 50, 75, 100, 140, 180 ms; a higher acceleration factor along *k_y_* = 3 for IR-GE and single-echo SE; and FOV of 3D-GRE = 216 × 176 × 210 mm^3^. Even with the lower resolution and higher acceleration, the total scan time was still about 53 minutes, including 27 minutes for single-echo SE, 14 minutes for IR-SE, and 12 minutes for GRE.

### 2.4. Image processing and analysis

Data reconstructions were performed in MATLAB using a Linux workstation (CPU: Intel Xeon, 3.00GHz, 24 Cores; RAM: 512 GB; GPU: Quadro RTX 5000, 16 GB memory). The subspace reconstruction was solved by the alternating direction method of multipliers (ADMM) algorithm (Boyd et al., 2011) implemented in the Berkeley Advanced Reconstruction Toolbox (BART) (Tamir et al., 2016; Uecker et al., 2015).

In the test-retest experiment, the quantitative maps from the two scans (3-min 1-mm protocol) were first registered using FSL FLIRT (Jenkinson et al., 2002; Jenkinson et al., 2012; Jenkinson and Smith, 2001). Then, the averaged R1 maps of each subject were used for Freesurfer (Desikan et al., 2006; Fischl, 2012; Fischl et al., 2002) segmentation, which resulted in 165 Region Of Interests (ROI) in cortical, subcortical, white matter and cerebellum regions, after removing CSF regions and ROIs smaller than 50 voxels. Two-sided *t*-tests and Pearson’s correlation coefficients were used in the repeatability analysis. Surface-based cortical reconstruction was performed using Freesurfer (Desikan et al., 2006; Fischl, 2012; Fischl et al., 2002) on the R1 maps separately for each subject and for each scan (9-min 0.7-mm protocol). 9 equi-volume (Waehnert et al., 2014; Waehnert et al., 2016) cortical layers were reconstructed, and applied to all quantitative maps to investigate their distribution across different cortical depths. These maps were sampled onto the average subject space with a 2-mm Gaussian surface smoothing for final analysis.

Synthetic images were synthesized from the calculated quantitative maps based on signal equations of specific sequences. To optimize visualization between two types of tissue of interest, a spectrum of contrast difference was obtained using a range of acquisition parameters. The mean value of the synthesized image intensity of the target tissues was used to calculate the contrast difference. The tissue can be identified by the acquired quantitative maps, for example, a threshold-hold based segmentation on T_1_ map was performed here to generate masks for gray matter, white matter and CSF.

## 3. Results

### 3.1. Simultaneous T_1_, T_2_, and T_2_* mapping in 3-minutes at 1-mm isotropic resolution

Figure 3 demonstrates an example dataset acquired by 3D-EPTI at 1-mm isotropic resolution with whole brain coverage in 3 minutes, resulting in a total of 1350 multi-contrast images resolved at a time interval as short as 0.93 ms (an echo-spacing). Representative reconstructed images with different T_1_, T_2_, T_2_* contrasts illustrate the high quality in the reconstruction that can be achieved from a highly undersampled dataset by fully exploiting the intra- and inter-readout spatiotemporal correlations (Fig. 3a). The resultant quantitative parameter T_1_, T_2_, T_2_*, PD maps show high image SNR and resolution, without image distortions and noticeable aliasing artifacts (Fig. 3b). In addition to these quantitative parameters, 3D-EPTI also estimates the B_1_^+^ field, therefore achieves improved accuracy for quantification without the need of an additional scan for B_1_^+^. These multi-parametric maps acquired in a single scan are also perfectly aligned without the need for co-registration. The accuracy of the quantitative estimates, including T_1_, T_2_, T_2_*, PD and B_1_^+^, was also tested through a simulation study with gold standard reference parameter maps, where low errors were observed for all parameters as shown in Supplementary Fig. 3.

**Figure 3.**
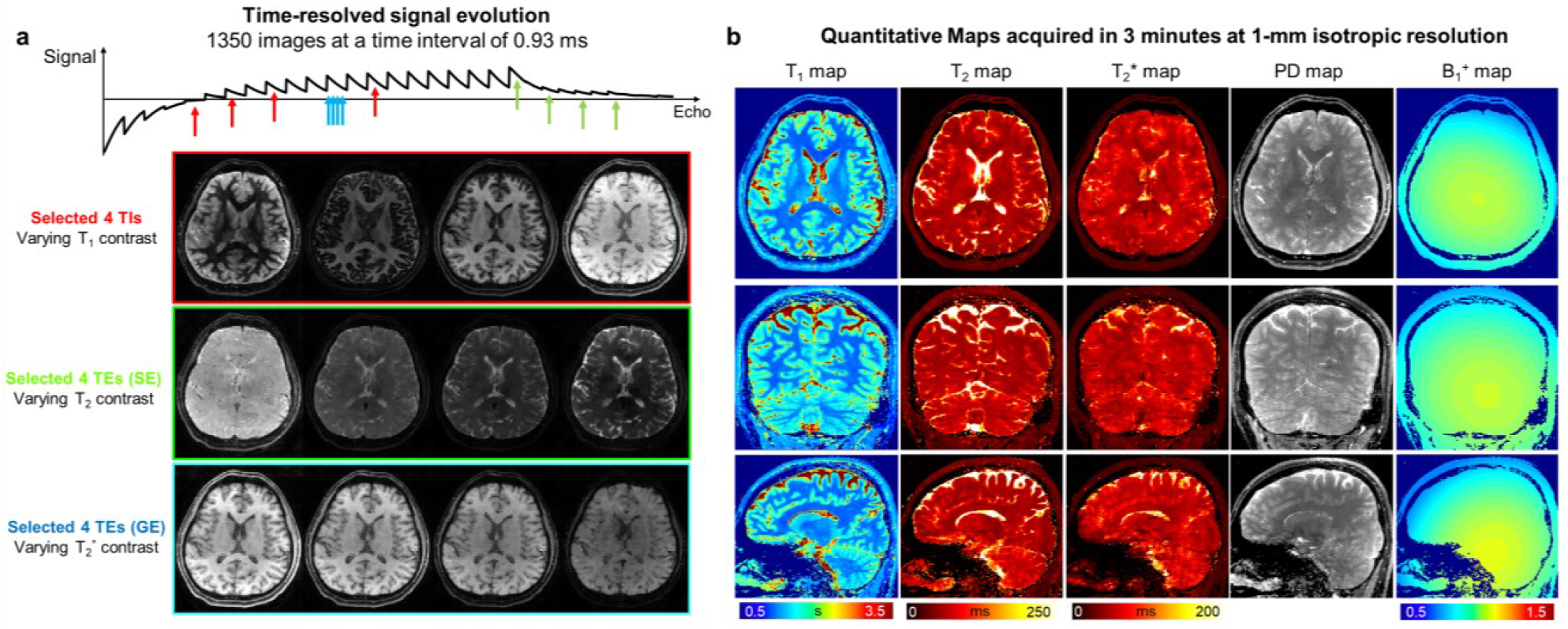
Simultaneous whole-brain T_1_, T_2_ and T_2_* mapping at 1-mm isotropic resolution in 3 minutes acquired by 3D-EPTI. **a**, Representative reconstructed multi-contrast images with different T_1_, T_2_, and T_2_* weightings selected from the 1350 resolved images. **b**, High-resolution quantitative maps estimated from the multi-contrast images, including T_1_, T_2_, T_2_*, PD and B_1_^+^, shown in three orthogonal views.

### 3.2. Characterization of the repeatability and reliability for T_1_, T_2_, and T_2_* mapping

Quantitative repeatability is crucial for monitoring and characterization of brain tissues over time and among populations. To evaluate the repeatability of 3D-EPTI, the scan-rescan assessment performed on 5 healthy volunteers using the 3-minute 1-mm protocol is shown in Fig. 4. An example of the auto-segmented Freesurfer ROIs used in the study is demonstrated in Fig. 4a. High correlations and small differences were measured between the two scans for all three parameters as shown in Fig. 4b and 4c. Specifically, the T_1_ values from the first and the second scans are highly correlated with a positive Pearson’s correlation coefficient (PCC) of 0.996 (P < 0.0001). Bland Altman analysis revealed no significant difference (P = 0.821, two-tailed *t*-test) between the two measurements. The T_2_ values of the two scans show a high positive correlation (PCC = 0.974, P < 0.0001) with a small difference of 0.28% (P = 0.017), and the T_2_* values also show a high positive correlation (PCC = 0.938, P < 0.0001) and a small difference of 0.80% (P < 0.0001). The scan-rescan difference maps (Fig. 4e) show a low level of differences and a relative homogeneous distribution in the gray and white matter areas of interest, while large differences are mainly observed in the CSF regions, which could be attributed to the high physiological noise in these regions.

**Figure 4.**
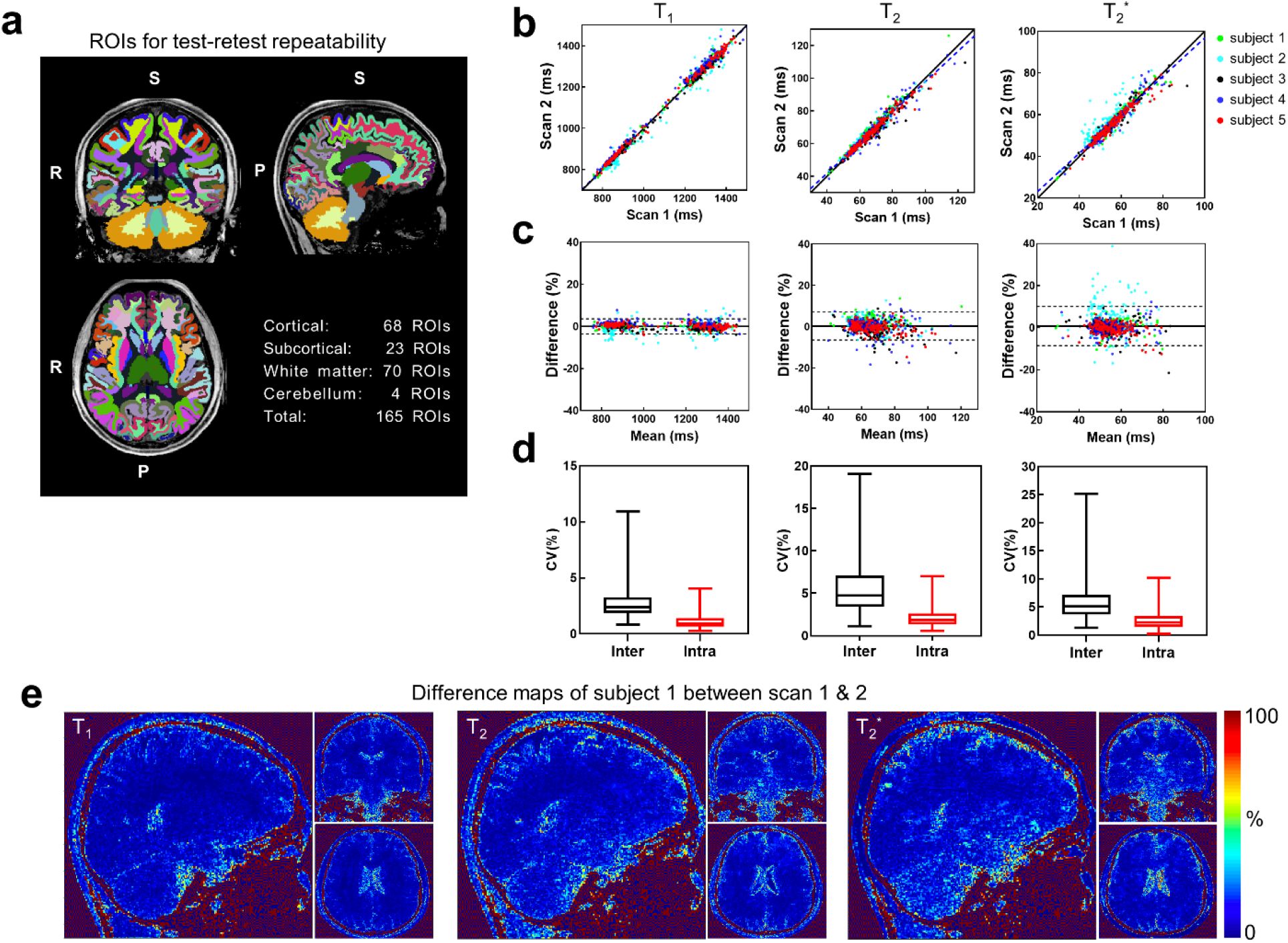
Repeatability test of 3D-EPTI for simultaneous T_1_, T_2_ and T_2_* mapping using the 3-minute 1-mm protocol. **a**, 3D volumes are segmented into 165 ROIs covering the whole-brain area. **b**, Scatter plots of the test-retest T_1_, T_2_, T_2_* values in the 165 ROIs measured from 5 subjects, shown along with the identity line (solid) and the regressed line (dashed). **c**, Bland-Altman plots of the test-retest quantitative parameters. Solid lines show the mean differences or the estimated biases, while the dotted lines show the 95% limits of agreements. **d**, Box-plots of the coefficient of variation (COV) between (inter) and within (intra) subjects with whiskers showing minimum and maximum data points. **e**, The percentage difference maps in one of the subjects that shows the spatial distribution of the test-retest variations for the 3 parameters.

The intra-subject variability between test-retest scans and inter-subject variability among the five volunteers were also evaluated and compared using coefficient of variation (COV). Figure 4d shows the distribution of the intra-subject and inter-subject COVs across all ROIs in box plots with whiskers from minimum to maximum data points. Small intra-subject variability was observed with a median COV at 0.93%, 1.88% and 2.27% for T_1_, T_2_ and T_2_* respectively. The inter-subject COVs were also measured to be at a low level, with median COVs of 2.39%, 4.75%, 5.09%, which are higher than the intra-subject measurements as expected. The overall low level of COVs demonstrates the high level of repeatability of the 3D-EPTI quantifications, while the expected differences between intra- and inter-subject suggest the potential ability of 3D-EPTI in capturing individual differences. The T_2_* values show the largest variability that could be reflective of the variability in the head position between the scan and rescan acquisitions, where previous findings have demonstrated variability in T_2_* values as a function of head orientation relative to the main field (Cohen-Adad et al., 2012). The repeatability of B_1_^+^ mapping was also evaluated where consistent measurements were obtained across three scans at different spatial resolutions for the same subject as shown in Supplementary Fig. 4.

To further validate 3D-EPTI’s reliability, a comparison between quantitative parameters provided by 3D-EPTI and those from standard acquisition methods was conducted *in vivo.* The well-established standard methods provide high quality quantitative parameters at a cost of impractically long acquisition time and therefore a higher level of susceptibility to motion induced artifacts. To mitigate this issue in our comparison, we reduced the spatial resolution (2 × 2 × 3 mm^3^ for T_1_ and T_2_) as well as the slice coverage of the standard acquisitions to keep them to an acceptable total acquisition time of 53 minutes. For 3D-EPTI, a single 3-minute scan at 1-mm isotropic resolution with whole brain coverage was used to obtain all the quantitative estimates. ROI analysis was performed in 14 manually selected ROIs that were contained within the slice coverage of the standard 2D acquisitions. As shown in Fig. 5, T_1_ measurements of 3D-EPTI and the standard method show high positive correlation (PCC = 0.972, P < 0.0001), with a bias of −7.23% (P < 0.0001). Similarly, T_2_ measurements also show high positive correlation (PCC = 0.790, P = 0.0008) with no significant bias (P = 0.1351). Lastly, T_2_* values are also highly correlated (PCC = 0.948, p < 0.0001) with a small bias of 4.68% (P = 0.0137). In addition to the in-vivo validation, a phantom experiment was also performed where the quantitative measurements from 3D-EPTI are in good agreement with the standard methods (Supplementary Fig. 5).

**Figure 5.**
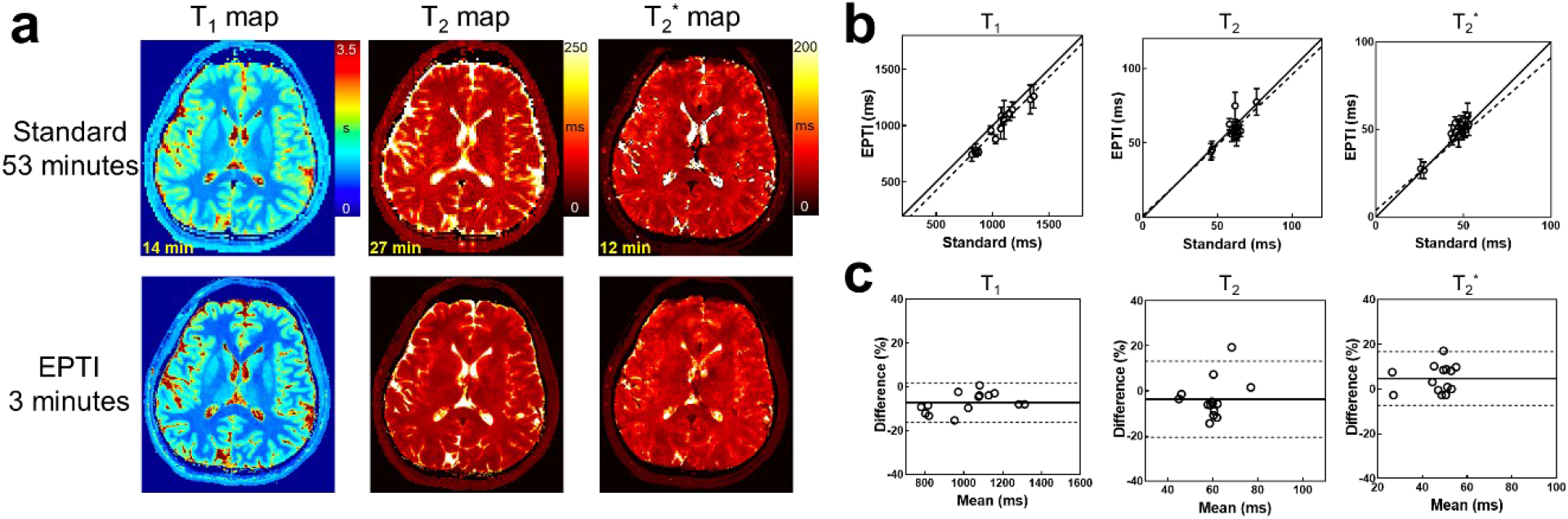
Comparison of the quantitative measurements obtained using 3D-EPTI vs. lengthy standard acquisitions *in vivo.* **a**, Quantitative maps acquired by the standard methods and 3D-EPTI. **b**, Scatter plots of the quantitative values from 14 selected ROIs, shown along with the identity line (solid) and the regressed line (dashed). **c**, Bland-Alman plots of the same data with the mean differences or the estimated biases (solid lines) and the 95% limits of agreements (dotted lines).

### 3.3. Ultra-fast 1-min scan and submillimeter mapping enabled by the high efficiency

In addition to the 3-minute 3D-EPTI protocol at 1-mm isotropic resolution, two additional whole-brain protocols at different resolutions were developed to showcase the high efficiency of 3D-EPTI.

First, an ultra-fast 1-minute protocol at 1.5-mm isotropic resolution was developed to obtain high quality quantitative maps as shown in Fig. 6 (T_1_ map) and Supplementary Fig. 6 (T_2_ and T_2_* maps). A one-minute ultra-fast scan has been an alluring target for the MRI research community, but to our knowledge, no study so far has been able to obtain whole brain quantitative parameters in a scan as short as 1 minute at a reasonable isotropic spatial resolution. The 1-minute multi-parametric scan provided by 3D-EPTI could help fulfill this important unmet need. The quick 3D-EPTI scan could potentially reduce the chance of involuntary movements, a major source of image artifacts in MRI, particularly in less compliant clinical patients (e.g., pediatric patients), as well as improve patient throughput and reduce costly re-scan, patients called backs (Andre et al., 2015) and the need of harmful anesthesia (Ing et al., 2012).

**Figure 6.**
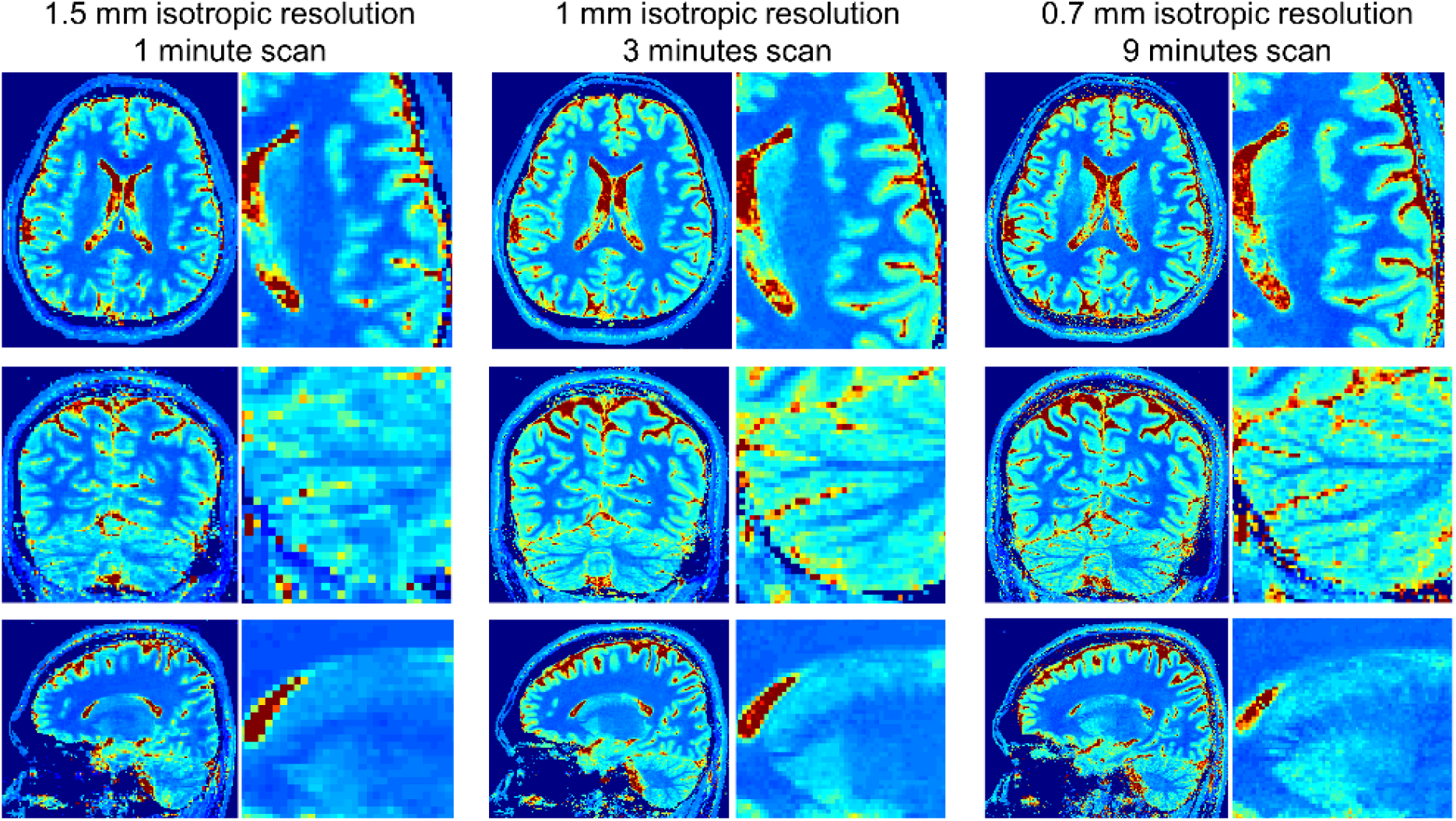
Example T_1_ maps with zoomed-in areas provided by 3D-EPTI protocols at different spatial resolutions: 1-minute scan at 1.5-mm isotropic resolution, 3-minute scan at 1-mm isotropic resolution, and 9-minutes scan at 0.7-mm isotropic resolution.

Another protocol has also been developed for the acquisition of 0.7-mm isotropic resolution quantitative parameters in 9 minutes, to allow visualization and analysis of more detailed brain structures (Fig. 6, right column). This should be particularly helpful in studying the intra-cortical architecture of the human brain, where the thin cortex consists of multiple layers with different tissue properties, such as different levels of myelination or iron concentration (Carey et al., 2018; Haast et al., 2016; Lutti et al., 2014; Marques et al., 2017; Trampel et al., 2019; Waehnert et al., 2016; Warntjes et al., 2016). The high efficiency of 3D-EPTI enables for the first-time simultaneous acquisition of T_1_, T_2_, T_2_* at submillimeter resolution within a few minutes, and provides a new powerful tool to study cortical layer-dependent tissue properties.

Here, we explored the feasibility of using 3D-EPTI to assess intra-cortical structures of healthy volunteers, evaluated its inter-scan and inter-subject repeatability, and explored its potential to identify specific spatial features. Figure 7a shows the quantitative maps of all three parameters at different cortical depth from inner (proximate to white matter) to outer (proximate to CSF) surface of a healthy volunteer. Consistent profiles across layers can be observed with distinctly lower values in areas that are highly myelinated, such as motor, sensory, auditory, MT and visual cortex, which are in accordance with previous literatures (Carey et al., 2018; Glasser et al., 2014; Haast et al., 2016; Lutti et al., 2014; Marques et al., 2017). For all three relaxation parameters, there is a general increasing trend across cortical layers from inner to outer surface, potentially reflecting the decrease of myelin and iron contents from the white matter surface to the CSF surface that has been validated in histological studies (Annese et al., 2004). Figure 7b shows an example of such profile within a representative ROI in the motor cortex (BA 4a), where consistent profiles from 4 healthy volunteers (color-coded) across two repeated scans (solid and dashed lines) were plotted, showing high inter-scan (intra-subject) and inter-subject repeatability. The high repeatability is also observed in all other representative ROIs as shown in Fig 7d and 7e, quantified by both the root mean square errors (RMSEs) and the PCC, which assess the differences and the similarities of the cortical profiles among subjects and between scans. The intra-subject profiles were measured with low differences (median RMSEs for T_1_, T_2_, T_2_* at 20.36 ms, 1.84 ms and 2.94 ms) and high correlations (median PCCs at 0.999, 0.999,0.992). The inter-subject profiles showed higher variability than intra-subject as expected, but still with low median RMSEs at 38.47 ms, 4.37 ms and 4.42 ms as well as high median PCCs at 0.998, 0.995, 0.988 for T_1_, T_2_, T_2_* respectively.

**Figure 7.**
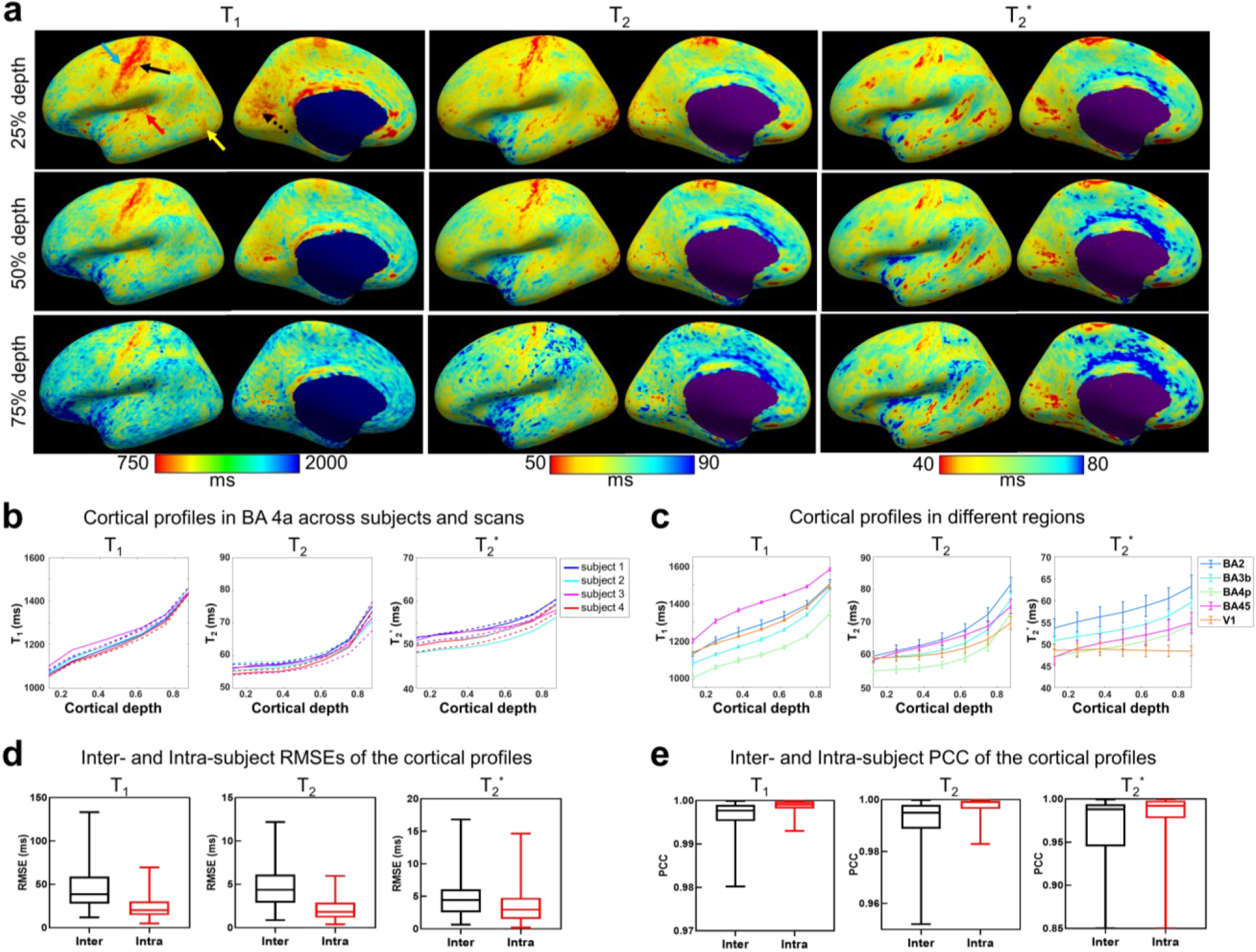
Surface-based cortical analysis of T_1_, T_2_ and T_2_* of 0.7-mm isotropic resolution 3D-EPTI data. **a**, Quantitative parameters sampled at three different cortical depths (25%, 50%, 75%) shown on the reconstructed cortical surface. Lower quantitative values were observed in highly-myelinated regions such as motor (blue arrow), somatosensory (black arrow), auditory (red arrow), middle temporal visual area (yellow arrow) and visual cortex (dotted black arrow). **b**, Example of quantitative values as a function of cortical depth in the BA 4a area of 4 subjects across 2 repeated scans (solid line: scan 1, dashed line: scan 2). **c**, Averaged quantitative values over the 2 scans across 4 subjects as a function of cortical depth in 5 representative ROIs. Error bars represent the standard deviations between the 4 subjects. **d**, Box plots of the root-mean-square-errors (RMSEs) of the cortical profiles of the quantitative parameters calculated between (inter) and within (intra) subjects. **e**, Box plots of Pearson correlation coefficients (PCCs) of the profiles calculated between (inter) and within (intra) subjects.

Despite the remarkable consistency across scans and subjects, the cortical profiles differ across different cortical regions. The average profiles in Fig. 7c revealed different slopes and global values in different ROIs. The results provided by 3D-EPTI in a dramatically reduced scan time agree well with previous literatures investigating T_1_ or T_2_* cortical profiles (Carey et al., 2018; Marques et al., 2017; Waehnert et al., 2016). For example, highly myelinated areas such as the motor (BA 4p), and the sensory (BA 3b) areas have globally lower T_1_ profiles than other areas. A relative flat T_2_* profile was observed in the visual cortex, which was also reported previously (Carey et al., 2018; Marques et al., 2017) and might be attributed to the highly myelinated middle layers and increasing susceptibility in the outer layers due to presence of blood vessels with a high level of iron. Previous studies investigating T_2_ profiles across cortical depths have been lacking due to the need for prohibitively long acquisitions. Using 3D-EPTI, different T_2_ profiles were also obtained in different ROIs, which could reflect differences in both myelin water content and iron composition and complement to the findings in T_1_ and T_2_*.

### 3.4. High quality synthetic images

Figure 8 demonstrated the contrast-synthesizing ability of 3D-EPTI using its co-registered quantitative maps in several examples, including the clinical routine contrasts (MPRAGE, T_2_-FLAIR and T_2_-weighted), and some advanced contrasts such as double inversion recovery (DIR) to provide superior cortical visualization (Calabrese et al., 2007; Wattjes et al., 2007). The high efficiency of 3D-EPTI allows us to achieve higher spatial resolution within clinical acceptable time, which significantly reduces the partial volume effects commonly observed in synthetic images (e.g., the 0.7-mm data shows minimal partial volume effects even in challenging FLAIR contrast compared to the 1-mm and 1.5-mm data). The high isotropic resolution of the 3D-EPTI protocols also renders it the flexibility of viewing the images in arbitrary orientations (Fig.8b). Using the quantitative parameters, the weighting of each of these contrasts is adjustable to provide optimal visualization for the tissue of interest without the need to re-scan the subject. For instance, the contrast-determining parameters of an IR-SE sequence (FLAIR), TI and TE, can be freely adjusted across a wide range, resulting in a spectrum of possible contrasts between tissues (Fig. 8c). From the spectrum, a particular set of parameters can be selected for final image synthesis, such as the ones that offer the maximum white-grey or CSF-gray contrast.

**Figure 8.**
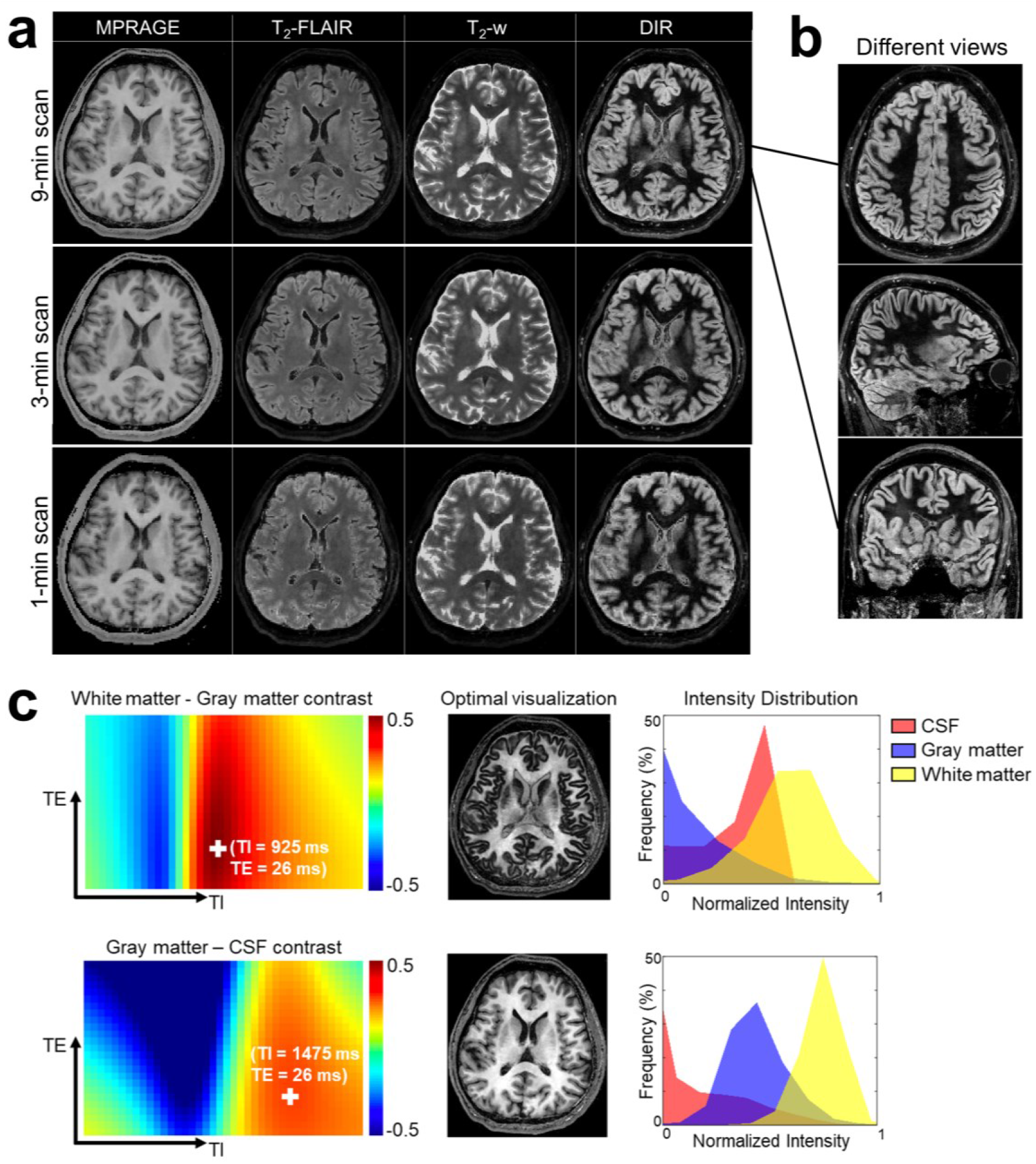
Synthesized multi-contrast images using 3D-EPTI. **a**, Synthetic MRRAGE, T_2_-FLAIR, T_2_-weighted, and double-inversion recovery (DIR) contrasts from three protocols (9-min 0.7-mm protocol, 3-min 1-mm protocol, and 1-min 1.5-mm protocol). **b**, An example of the flexibility in visualization in different views of the synthesized images at high isotropic resolution. **c**, Examples of contrast-optimized visualization between target-tissue pair by adjusting sequence parameters (TI and TE). Two examples that maximize the contrast differences i) between white and gray matter, and ii) between gray matter and CSF, are presented. The spectrum of contrast difference between the two tissues (left panel) is used to select the parameters to synthesize the contrast for optimal visualization (middle panel). The intensity distribution of the optimized contrast in three tissues are shown on the right.

## 4. Discussion

The goal of this study is to address the long-standing problem in quantitative MRI — the slow acquisition speed. The key concept in 3D-EPTI of exploiting spatiotemporal correlation at multiple timescales through new encoding strategies within and between its efficient continuous readouts was used to achieve rapid quantitative MRI. This has allowed the acquisition of robust multi-parametric maps within minutes at high isotropic resolution with whole brain coverage. As a proof-of-concept, a 3D-EPTI sequence for simultaneous T_1_, T_2_ and T_2_* mapping was developed and validated.

High intra-subject repeatability of the quantitative parameters obtained using 3D-EPTI was demonstrated across multiple healthy volunteers. This will be critical to the success of future deployment of 3D-EPTI to various applications such as in longitudinal monitoring of healthy and diseased tissues during complex biological process of brain development or pathophysiological progression of neurological diseases. Nonetheless, despite our efforts to minimize the bias in the repeatability assessment process itself, such as by using automatically segmented ROIs instead of manual ROIs, the small variations (bias < 0.8% and COV < 2.27%) could still be partially caused by errors in the registration process to align the two scans for comparison. Moreover, inherent differences between scans are possible. For example, T_2_* shows slightly higher variability than T_1_ and T_2_, which may be explained by the differences due to its dependence on head orientations relative to the main magnetic field.

The high inter-subject repeatability (COV < 5.09%) of the quantitative parameters obtained using 3D-EPTI on healthy volunteers points to its potential for use in establishing population-average norms or atlases. Currently, multi-parametric MRI atlases are still lacking, but the short scans enabled by 3D-EPTI can significantly improve the cost effectiveness and facilitate large-scale studies for this purpose. On the other hand, as expected, the inter-subject repeatability is lower than the intra-subject repeatability. This could reflect the ability of 3D-EPTI to detect inherent individual differences, pointing to its potential in providing sensitive quantitative biomarkers. However, some of the increased variability could also be attributed to the additional segmentation variabilities across subjects, especially in challenging areas. High intra- and inter-subject repeatability was also observed in the intra-cortical profiles obtained by 3D-EPTI. This illustrates the robust performance in using 3D-EPTI data to conduct reliable surface reconstruction and reveal repeatable subtle features across cortical layers. In addition to its potential for use in longitudinal monitoring and in establishing quantitative biomarkers, 3D-EPTI could be used to evaluate spatially varying profiles of cortical myelination and iron concentration.

The reliability of the quantitative measures obtained using 3D-EPTI was further validated by comparing them to ones obtained using lengthy standard acquisitions, where a generally high level of agreement was observed in both in-vivo (Fig. 5) and phantom (Supplementary Fig. 5) experiments. The largest difference was observed in the in-vivo T_1_-values, with a bias level of 7.23%. This could potentially be attributed to the magnetic transfer (MT) effect, which causes a different level of exchange between water and macromolecular pools in different sequences. This can create discrepancies between the actual and the modelled signal evolutions in both 3D-EPTI and in the standard acquisition, where the MT effect has not been accounted for. Such an effect is more prominent in T_1_ quantification, due to the use of an inversion recovery pulse. The characterization and disentanglement of the MT effect in quantitative mapping is an active area of research (Hilbert et al., 2020). Future work will investigate the incorporation of a MT-sensitive sequence module into the 3D-EPTI acquisition, and model the MT effect in both reconstruction and parameter fitting.

The concept of 3D-EPTI can be readily adopted to other pulse sequences for the quantification of other tissue parameters. For example, 3D-EPTI should be exceptionally well suited for parameters estimation for multiple-pool models (Dong et al., 2021a), because it can continuously track complex signal evolution at a very short time scale (< 1 ms) to offer more degrees of freedom in estimating these parameters. The unique features of 3D-EPTI also open up many possibilities for further technical improvements. For example, the radial-block sampling of 3D-EPTI grants it the ability to monitor and correct subject movements between repetitions for motion-robust scanning (Dong et al., 2021b). Other advanced reconstruction algorithms, including machine learning and multi-dimensional low-rank tensor approach, may further improve the accuracy of time-resolving reconstruction. The adaptation of EPTI to non-Cartesian acquisitions can also help eliminate their potential distortion/blurring artifacts due to the long readouts and provide multi-contrast capability (Fair et al., 2020; Liberman et al., 2020a, b).

To summarize, we introduce a novel method, termed 3D-Echo Planar Time-Resolved Imaging (3D-EPTI), that enables high acceleration and significantly improved imaging efficiency for multi-parametric imaging. By pushing multi-parametric MRI into a fast regime (as short as 1 minute), 3D-EPTI should help facilitate a paradigm shift from qualitative to quantitative imaging in clinical practice, and also open up the possibility of acquiring multi-parametric maps at submillimeter isotropic resolution within a few minutes to reveal exquisite brain structures for neuroscientific research. With richer and more reproducible information obtained within a single scan, 3D-EPTI has the potential to increase patient throughput, patient compliance and cost effectiveness, paving the way for large-scale studies to establish new quantitative biomarkers for neurological diseases.

## Acknowledgments

This work was supported by NIH NIBIB (R01-EB020613, R01-EB019437, R01-MH116173, P41-EB030006 and U01-EB025162) and by the MGH/HST Athinoula A. Martinos Center for Biomedical Imaging; and was made possible by the resources provided by NIH Shared Instrumentation Grants S10-RR023401, S10-RR023043, and S10-RR019307. We also thank Dr. Gliad Liberman’s help on the GPU implementation of the image reconstruction.

## Supplementary Information

**Supplementary Figure 1.**
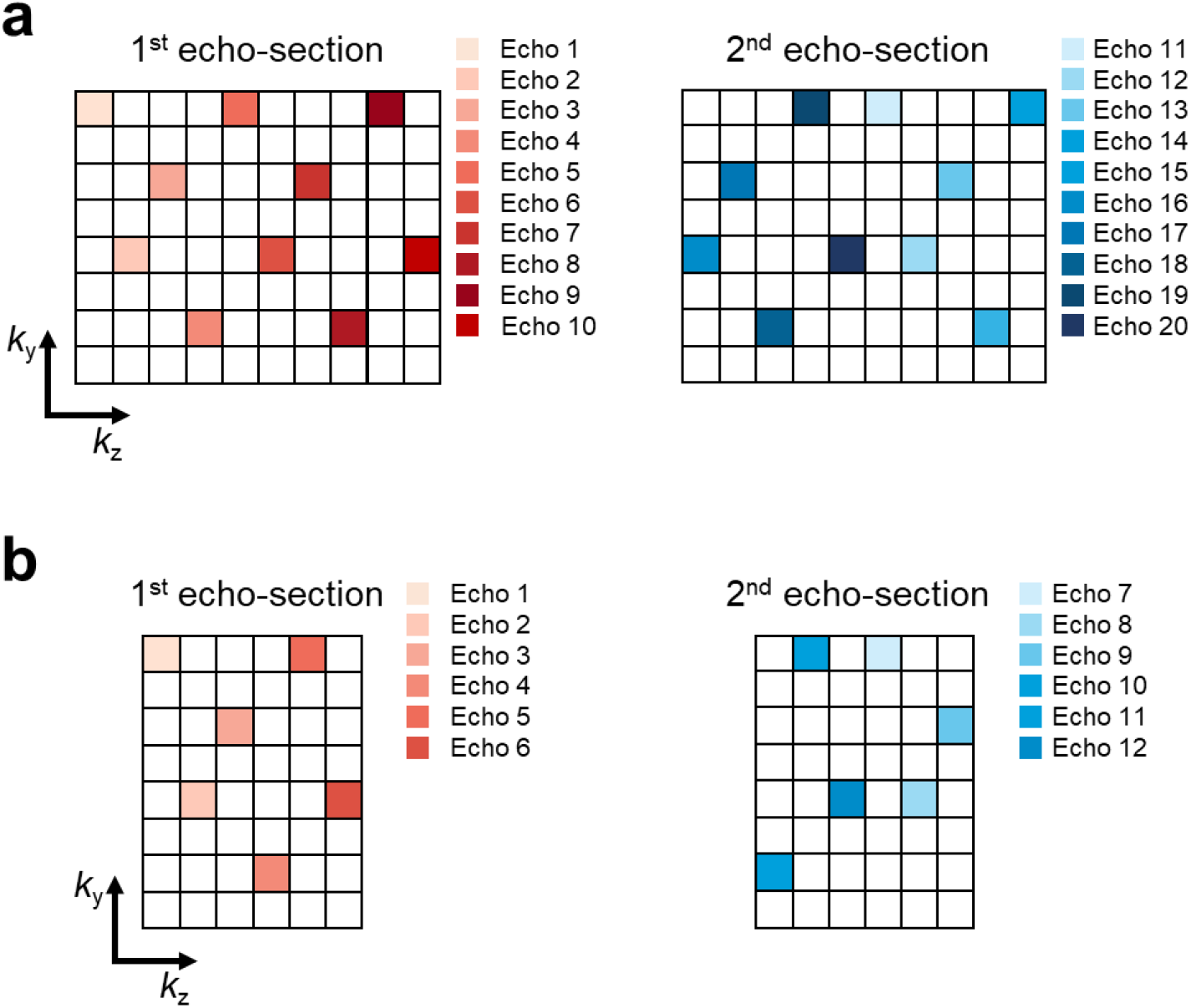
The encoding patterns in *k_y_-k_z_-t* space for (a) the 1.5-mm and 1-mm protocols, and (b) the 0.7 mm protocol. Two CAIPI patterns (red and blue) are interleaved across echoes in different echo sections to provide more complementary samplings (i.e., red for 1^st^ echo section, blue for 2^nd^, red for 3^rd^, blue for 4^th^, etc.). An acceleration factor of 80 (*k_y_ × k_z_* = 8 × 10) was used for the 1.5-mm and 1-mm acquisitions, while a smaller acceleration factor of 48 (*k_y_ × k_z_* = 8 × 6) was used for the 0.7-mm acquisition to compensate for the larger time interval (larger echo spacing).

**Supplementary Figure 2.**
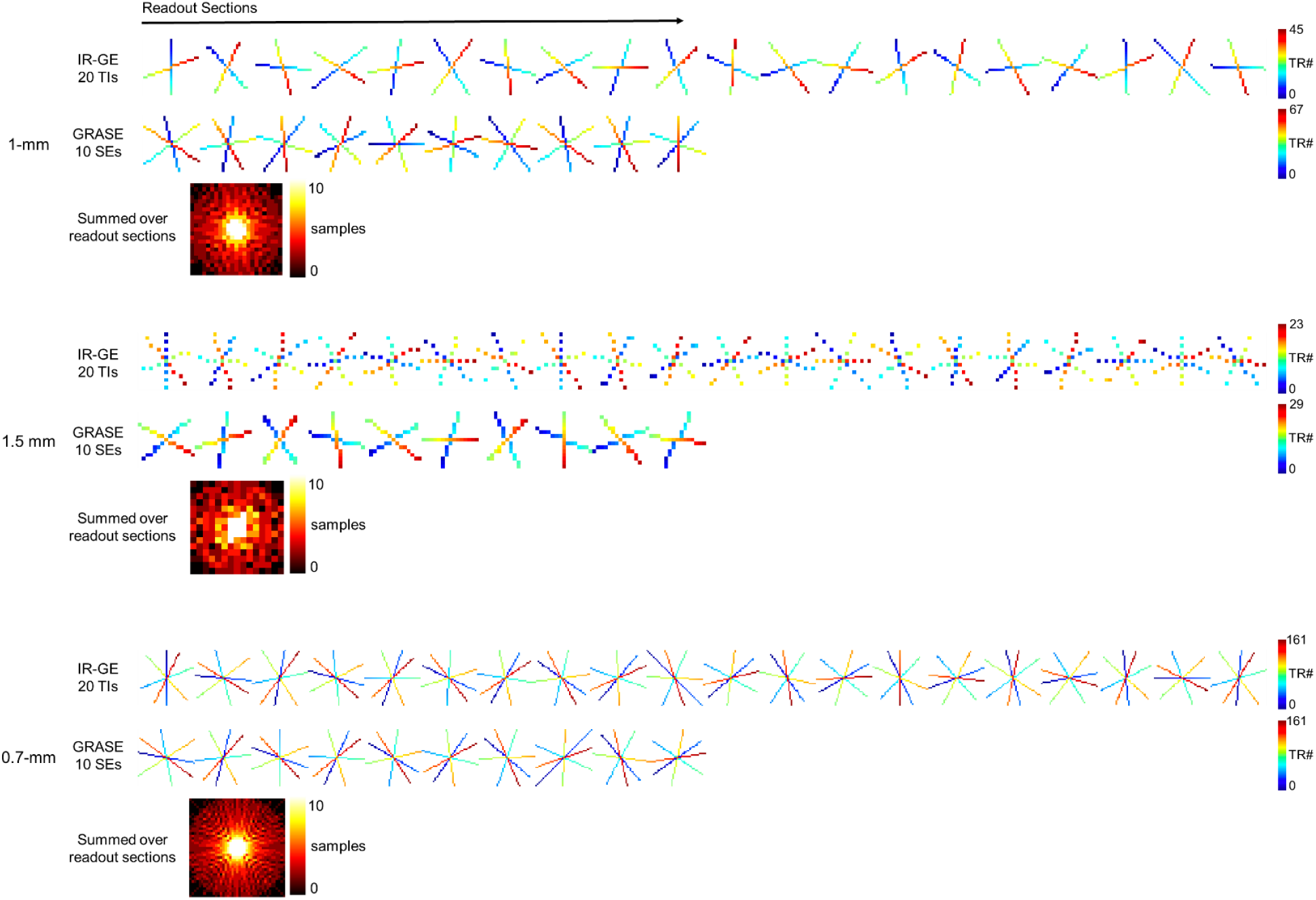
The golden-angle radial-block sampling pattern used in the designed protocols (top: 1-mm protocol, middle: 1.5-mm protocol, bottom: 0.7-mm protocol). Each color-coded point represents one sampling block acquired in a 3D-EPTI readout. After a few TRs, the sampling blocks acquired in the same readout section will form a radial-block pattern. The color coding represents the acquisition order of the blocks. For example, the index of TR (TR#) counts from 0 to the maximum, illustrated from blue to red. The combined sampling patterns across all readout sections in IR-GE and VFA-GRASE result in a variable density pattern as shown in the bottom.

**Supplementary Figure 3.**
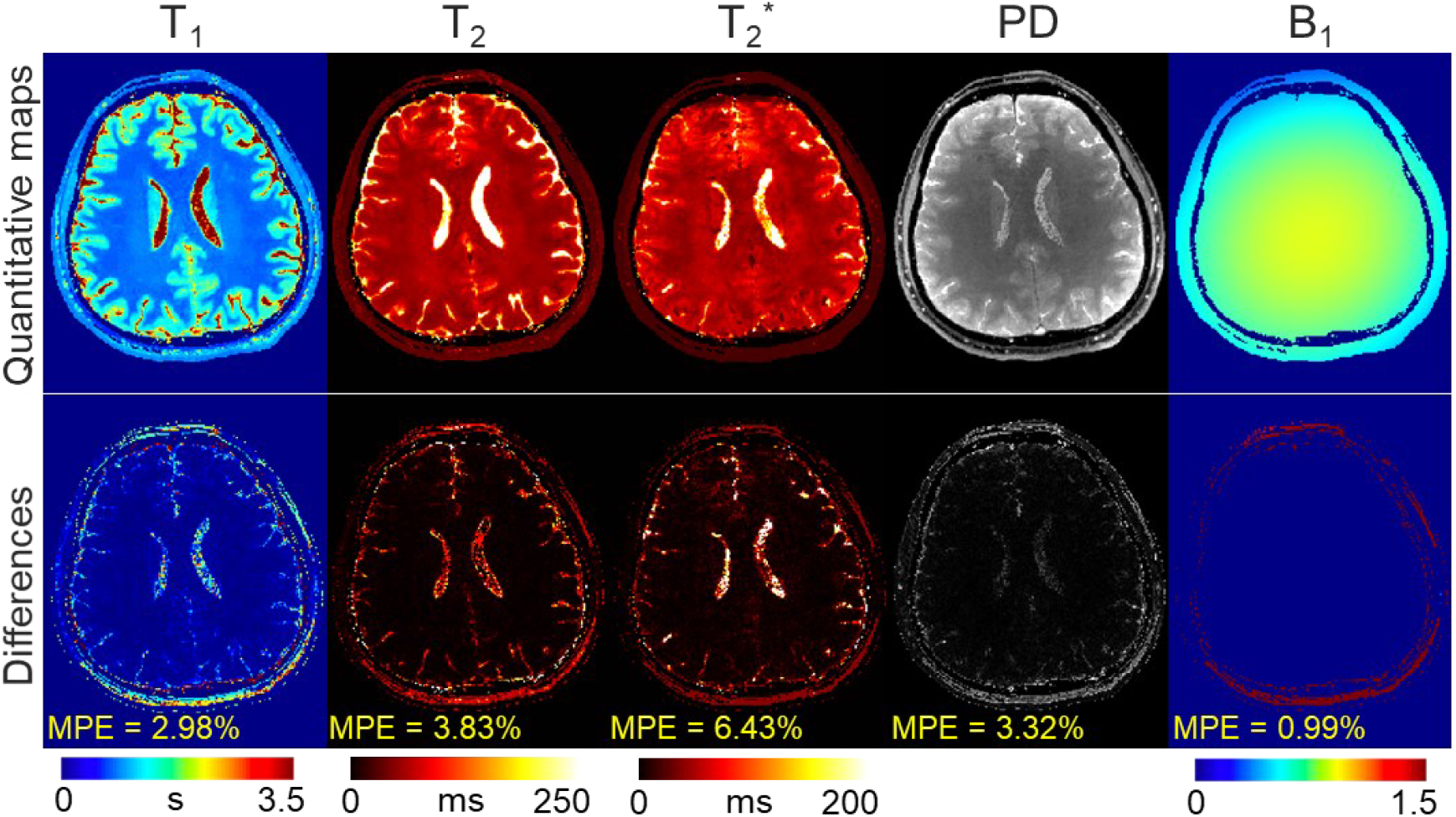
Results of the simulation experiment. The quantitative maps obtained by 3D-EPTI (3-min 1-mm protocol) were compared to the gold standard reference maps that were used to simulate the 3D-EPTI data. The difference maps (×2: magnified by a factor of 2) were calculated by subtracting the reference maps from the estimated quantitative maps. MPE: mean percentage error.

**Supplementary Figure 4.**
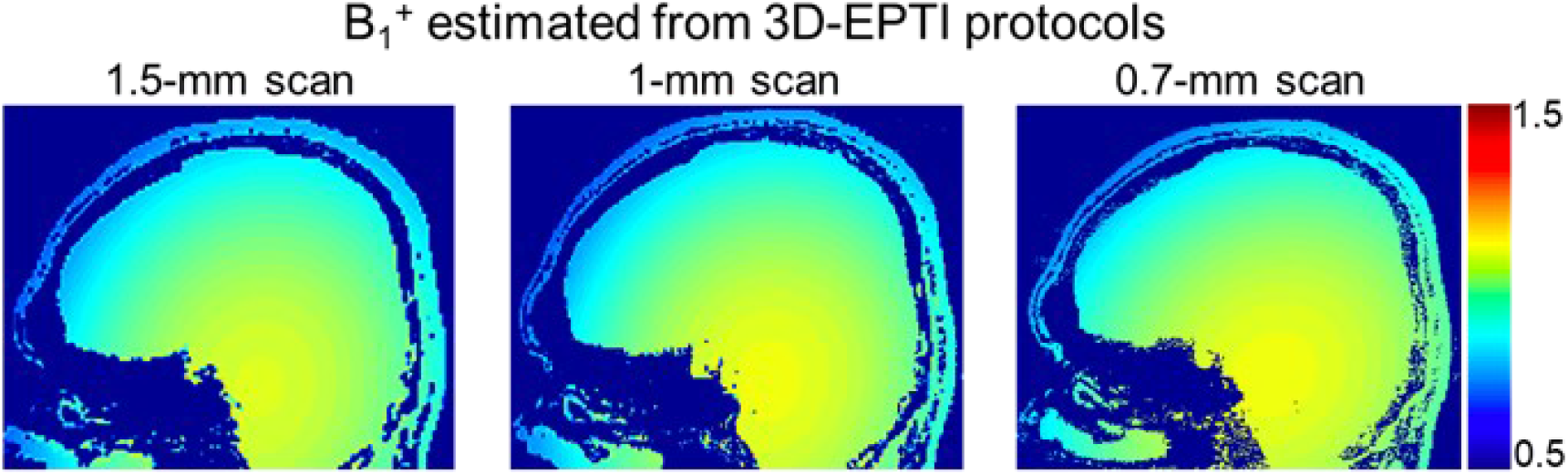
The estimation of B_1_^+^ field on the same subject from three 3D-EPTI scans at different resolutions.

**Supplementary Figure 5.**
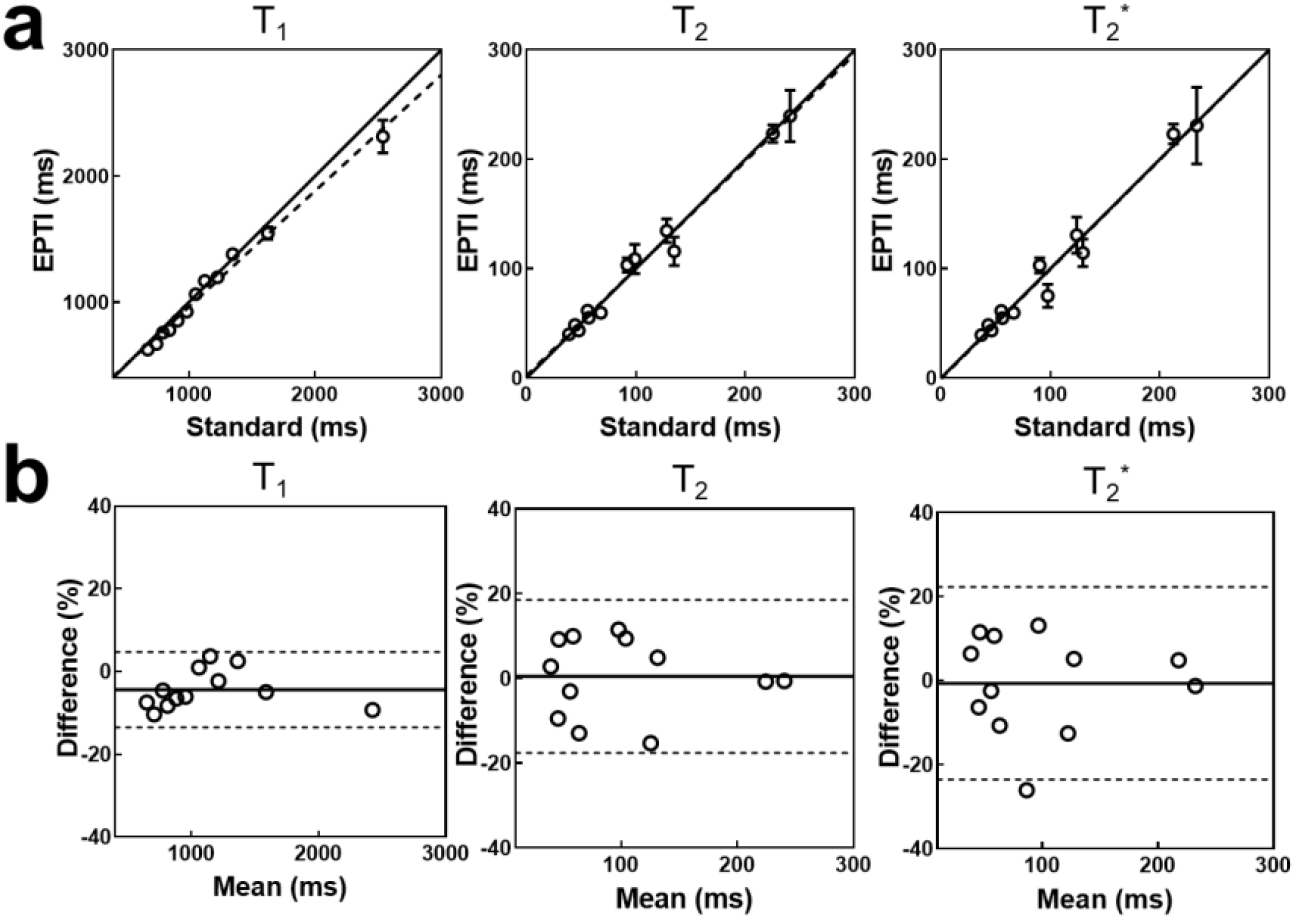
Comparison of the quantitative measurements between 3D-EPTI and standard acquisitions in the phantom experiment. **a**, Scatter plots of the quantitative values in 12 ROIs, shown along with the identity line (solid) and the regressed line (dashed). **b**, Bland-Alman plots of the same data with the mean differences or the estimated biases (solid lines) and the 95% limits of agreements (dotted lines).

**Supplementary Figure 6.**
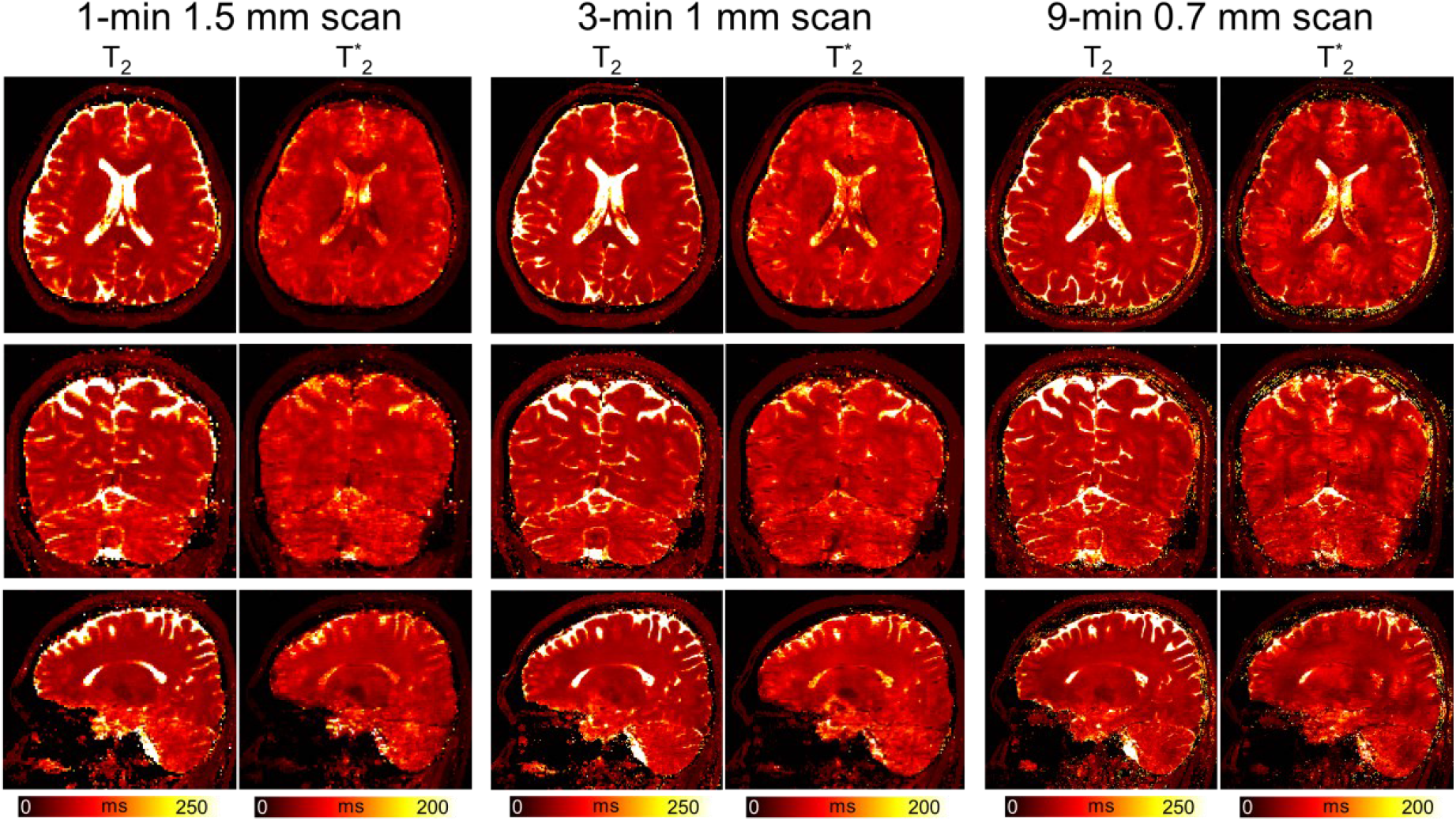
The estimated T_2_ and T_2_* maps from three different protocols on the same subject. Left panel: 1-minute protocol at 1.5-mm isotropic resolution; middle panel: 3-minute protocol at 1-mm isotropic resolution; right panel: 9-minute protocol at 0.7-mm isotropic resolution.

